# Arrayed hydrogels pair whole-cell imaging with single-cell mass spectrometry proteomics

**DOI:** 10.64898/2026.09.04.749469

**Authors:** Maya M. Overton, Cyril Deroy, Eileen Wang, Amber Lennon, Joshua E. Elias, Ryan McClure, Amy E. Herr

**Affiliations:** Department of Bioengineering, University of California, Berkeley, 342 Stanley Hall, Berkeley, CA 94720, United States; Biohub San Francisco, 499 Illinois St, San Francisco, CA 94158; Biohub Chicago, CZBiohub Chicago, 400 N Aberdeen St STE 800, Chicago, IL 60642, United States

## Abstract

Multimodal single-cell analysis aims to elucidate cellular-level phenotype within heterogeneous cell populations. Advancements in single-cell proteomics (SCP) seek to deepen quantitative depth, coverage, and reproducibility for a robust view of functional cell state. However, broadly accessible multimodal approaches are poised to benefit from sample-preparation advancements upstream of the mass spectrometer. Here, we introduce ProteoParcel, a multimodal SCP platform for indexing upstream widefield single-cell images to downstream label-free, bottom-up proteomics. Parcels are spatially arrayed planar polyacrylamide gels patterned with microwells. Each parcel – containing one microwell and an abutting gel region – is designed to integrate the single-cell imaging and SCP analysis modes. First, for whole-cell imaging, each microwell isolates an intact, individual breast cancer cell (MCF-7). After imaging, cells are subjected to in-microwell chemical cell lysis, electro-injection of whole-cell lysate from the microwell into the abutting gel region, in-gel chemical fixation, and finally in-gel tryptic digestion prior to peptide extraction for SCP. Location-indexed parcels are independently releasable to confer single-cell resolution to downstream mass spectrometry. To ensure SCP-suitable proteome solubilization and trypsin/Lys-C digestion, we optimize cell lysis, electrophoresis, and gel pre-equilibration conditions within the gel. Using ProteoParcel, we identify over 1,400 protein species from single, imaged MCF-7 cells. Scrutiny of the gel preparation conditions confirms that hydrogel-lysate interactions introduce predictable, physicochemically interpretable (cell membrane, hydrophobicity) detection biases, while broad functional-class composition and subcellular compartment coverage are preserved when benchmarked to in-solution digestion. ProteoParcel makes facile same-cell multimodal SCP and live-cell imaging.

## INTRODUCTION

Mass spectrometry (MS)-based proteomics identifies protein-level signatures of diseases like cancer and neurodegenerative diseases that remain undetectable through the lens of transcriptomic or genomic analysis alone.^1^ In diseased tissue with highly heterogeneous cell populations, single-cell resolution measurements detect divergent or rare-cell subpopulations defined by unique biomolecular signatures that are otherwise masked in bulk measurements. While single-cell RNA-sequencing approaches have transformed our understanding of gene expression heterogeneity, mRNA abundance is an imperfect proxy for protein content.^2^ Single proteomes, representing the active effector phenotype, are stable reflections of cellular state that can be interpreted deterministically across the analyte dynamic range.^3^ Single-cell proteomics (SCP) therefore annotates active functional cell states and supports discovery of unique biomarkers of disease and progression.^4–6^ Despite the power of SCP, multimodal integration provides an additional axis for decoding the relationship between proteome and phenotype.

Most existing multimodal SCP approaches are limited to targeted, antibody-based readouts^7–9^ or require highly specialized equipment and operators for execution, particularly in MS-based approaches.^10^ While ongoing research seeks to expand SCP accessibility through simpler workflows without sacrificing quantitative depth,^11^ pairing live-cell imaging with deep quantitative proteomics has been identified as a key gap in same-cell assays.^12^ The closest existing technology pairing imaging with MS-based proteomics, Deep Visual Proteomics (DVP), combines AI-driven image analysis with automated laser microdissection and ultra-high-sensitivity MS, but operates on fixed or archived tissue sections rather than live.^13^

Same-cell multimodal measurements depend on platforms that can isolate and spatially index individual cells while preserving cell identity when information is transferred off-chip. Many conventional approaches rely on microfluidic droplets for cell isolation and molecular indexing (i.e., barcoding). Droplet workflows isolate single cells in aqueous compartments and frequently co-encapsulate a barcoded bead that indexes each cell’s nucleic acids for pooled sequencing.^14,15^ The same architecture can also carry protein readouts: CITE-seq labels cells with barcoded antibodies to recover surface proteins and the transcriptome from one cell,^7^ and suspendable hydrogel nanovials pair secreted-protein capture with imaging, sorting, and downstream sequencing.^16^ Pairing droplets with MS has been a more difficult challenge because the carrier oil interferes with LC-MS/MS and must be removed. A recent approach uses digital microfluidics to cast each digest in an agarose droplet that solidifies and separates from the oil for transfer to an LC-MS/MS vial.^17^ Two constraints persist across droplet approaches: (1) cells partition into droplets by Poisson statistics rather than by deterministic placement, so single-cell occupancy is probabilistic; (2) the water-in-oil compartment also holds each cell behind a moving, curved interface in a bulk emulsion, a geometry ill-suited to time-resolved imaging of an individually addressed cell or to manual recovery of a chosen sample.

Planar arrays of microfabricated structures have been developed to address sample handling constraints through physically releasable substrates that preserve cell identity through downstream processing. In the microraft system, individual cells are isolated on detachable polymer elements arrayed on a compliant polydimethylsiloxane substrate.^18,19^ Target cells are identified by brightfield or fluorescence imaging, released by needle actuation, and collected for downstream culturing and clonal expansion. Within our own group, a related concept was applied to single-cell protein analysis.^20^ Spatially isolated polyacrylamide gel (PAG) elements patterned with microwells were used for single-cell western blotting (scWB), with the releasable particle format improving immunoprobing efficiency relative to previously established methods with on-chip readout.^21^ PAGE has long been used for targeted analysis of protein expression and purification of proteins by molecular mass for subsequent LC-MS/MS analysis.^22–24^ Together, spatially indexed, releasable PAG substrates can preserve cell identity through downstream molecular measurement, motivating the adaptation of PAGE with in-gel digestion as a sample preparation scaffold for LC-MS/MS in the single-cell regime.

The convergence of these capabilities – spatially indexed cell isolation, releasable gel-based substrates, and encapsulated, in-gel sample preparation – motivates ProteoParcel.

Fundamentally, ProteoParcel is a tool to make SCP compatible with live-cell imaging. ProteoParcel’s hallmark functionality arises from arrayed mesoscale PAG “parcels”, each of which is stippled with a 60-µm diameter microwell. Brightfield imaging verifies single-cell occupancy and indexes morphology (i.e., cell diameter, surface area) to downstream SCP. Cellular lysate is transferred from the microwell into the hygrogel parcel via electro-injection and fixation, and subsequently each parcel is released to a well-plate for in-gel digestion and peptide extraction for LC-MS/MS. The device supports manual manipulation of single proteomes for transfer of cellular materials from microscale structures to macroscale 384-well plates. Here we determined the proteomic depth and reproducibility of this approach for single cells and small populations of cells, as compared to a one-pot solution-based LC-MS sample preparation. Importantly, we characterize how ProteoParcel affects the physicochemical composition of the detected proteome by annotating proteins from two distinct gel preparation methods that differ in key electrophoresis and digestion chemistries.

## EXPERIMENTAL SECTION

### Device Fabrication

Negative molds were fabricated on silicon wafers using standard photolithography.^25^ SU-8 features were patterned at a height of 125 µm. Each SU-8 micropost forming the PAG microwell was 60 µm (optimal condition) or 100 µm (baseline condition) in diameter, and each SU-8 parcel mold was 2 mm × 5 mm. The patterned wafer was exposed to an O_2_/water vapor plasma at 100 W for 20 s. Immediately after plasma treatment, 800 µL of PAG precursor (7% T, 29:1) with 4% Rhinohide PAG Strengthener Concentrate (R33400, Invitrogen) was added atop the wafer and spread across all individual parcel molds using a P1000 pipette tip. A glass slide functionalized with 3-(trimethoxysilyl)propyl methacrylate was gently placed on the wafer and pressed down to ensure contact with the SU-8 features.^25^ The PAG precursor was polymerized for 20 min before the assembly was submerged in water for 30 min to hydrate the polymerized gel. The glass slide was removed from the wafer using a razor blade and the patterned ProteoParcel device was washed in deionized water for 10 min before drying with a stream of N_2_.

### Cell Culture

MCF-7/GFP cells were cultured in 1✕ RPMI Medium 1640 with L-glutamine (11875-093, Gibco) supplemented with 1% v/v penicillin streptomycin (15140-122, Gibco) and 10% BenchMark Fetal Bovine Serum (100-106, GeminiBio) in an incubator at 37 °C with humidified 5% CO_2_ air.

### Automated Cell Loading

Cell loading was performed as previously described.^26^ Adherent MCF-7/GFP cells were released by incubation in 0.05% trypsin-EDTA (25300-054, Gibco). Cells were washed twice with PBS before resuspension to 200,000 cells/mL in PBS. The cellenONE X1 (Cellenion) cell sorting and liquid handling system was primed according to the manufacturer’s instructions. The ProteoParcel device was manually aligned in the cellenONE to ensure the microwell array is parallel to the axis of motion of the liquid handler. The Piezo Dispense Capillary (PDC) was loaded with 5-10 µL of the MCF-7 cell suspension, and the appropriate number of cells were dispensed into each microwell (1-10 cells per microwell). The detection gating parameters for cell loading were a minimum cell diameter of 10 µm, maximum cell diameter of 100 µm, and elongation factor (length to width) of 5. The isolation gating parameters for cell loading were a minimum cell diameter of 18 µm, maximum cell diameter of 28 µm, and elongation factor of 3. Before removing the device, the brightfield images of each cell suspended in the cellenONE PDC were checked for doublets or abnormalities, and any parcels associated with upstream, intact-cell abnormalities were not analyzed.

### Imaging and Protein Staining

Brightfield imaging was performed after cells were seated in each microwell to verify cell loading and extract morphology (i.e., cell diameter, perimeter, area). The ProteoParcel device, a dried gel array fabricated on a 25 mm × 75 mm microscope slide, was imaged using an Olympus IX83 inverted microscope with a Prime BSI Express sCMOS camera (1T-01-PRIME-BSI-EXP, Photometrics). Images of the device were acquired at room temperature using a 40× objective. Cell diameter and area were extracted using ImageJ (ImageJ 1.53k java 13.0.6, NIH).

### Electrophoresis and Fixation

After cell deposition and optional brightfield imaging with the IX83 system, the devices prepared under the optimal ProteoParcel workflow were rehydrated in PBS for 5 min prior to being placed in a 60 mm × 100 mm horizontal electrophoresis chamber. Baseline ProteoParcel conditions were immediately subjected to a lysis buffer in the electrophoresis chamber after cell deposition. Cells were lysed (30 s) in a dual lysis/electrophoresis buffer (baseline: 0.5% SDS, 0.25% sodium deoxycholate, 0.5× Tris-glycine, 0.1% Triton™ X-100; optimal: 1% SDS, 0.5% sodium deoxycholate, 0.5× Tris-glycine, 0.2% Triton™ X-100), and proteins were electrophoresed into the gel at 40 V/cm for 15 s (baseline condition) or 20 s (optimal condition). Devices were removed from the electrophoresis chamber, placed in a 4-well dish, and fixed in a solution of 5% acetic acid and 50% methanol for 30 min on a rotator.

### Protein Staining

Total protein staining was performed using SYPRO™ Ruby Protein Gel Stain (S12000, ThermoFisher). After lysis, electrophoresis, and fixation, the device was transferred to a 4-well dish and covered with SYPRO™ Ruby. The device was heated to 80°C in a microwave with gentle agitation every 3 s. The device was stained for 5 min on a rotator before re-heating to 80°C in a microwave followed by a 25 min incubation on a rotator. The device was transferred to a separate well and washed with a 10% methanol and 7% acetic acid solution for 30 min on a rotator followed by three 5 min washes in UltraPure™ Distilled Water (10977015, ThermoFisher). The device was imaged on the GenePix 4300A microarray scanner (Molecular Devices) for SYPRO™ Ruby fluorescence with the 488-filter set at a resolution of 5 µm/pixel.

### Sample Preparation

Select parcels were removed from the glass slide substrate using a razor blade and transferred manually using sterilized tweezers to a 384-well plate containing 20 µL 100 mM ammonium bicarbonate (ABC) per well. After transfer of all parcels, the plate was centrifuged at 1500 g for 1 min and placed on a rotator at 200 RPM for 20 min. An equal volume of 100% acetonitrile (ACN) was added to each well, mixed by pipetting, and placed on a rotator for 20 min. Samples were dried in a SpeedVac vacuum concentrator at medium setting for 1 hr. In-gel digestion was prepared by addition of 10 µL of Rapid-Digestion mix: 2 ng trypsin/Lys-C (VA1061, Promega) with 0.05% *n*-Dodecyl β-D-maltoside (D4641-500MG, Sigma-Aldrich) in Rapid Digestion Buffer (VA1061, Promega) followed by centrifugation at 1500 g for 1 min. The plate was placed on ice for 1 h to allow the digestion mix to diffuse into the parcels. In-gel digestion was performed at 70°C for 1 h (baseline condition) or 2 h (optimal condition) in a HybEZ™ II Hybridization System (ACD Biotechne) containing deionized water-soaked

Kimtech™ Kimwipes™ to minimize evaporation. After incubation, plates were cooled to 4°C and centrifuged at 1500 g for 1 min. Formic acid (FA) and ACN were added to each well to a final concentration of 1% FA and 5% ACN and incubated on a rotator for 20 min for peptide solubilization. An equal volume of 100% ACN was added to each well for osmotic peptide extraction. Each liquid digest was transferred to an adjacent well and parcels were subjected to solubilization and osmotic extraction again before pooling the liquid digests. The parcels remained in the original well and were not transferred for LC. Plates were frozen at -80°C until LC loading.

### LC Loading

ProteoParcel digests were briefly dried in a Centrivap (LabConco Corporation, Kansas City, MO, USA) to remove residual acetonitrile. Samples were then resuspended in 30 µL of 0.1% FA, 0.015% DDM in water and pipetted repeatedly to ensure adequate resuspension prior to loading. Each sample was applied onto Evotip Pure C18 trap columns (Evotip, Evosep Biosystems, Denmark) following the manufacturer’s protocol. Evotips were conditioned with 20 µL of 0.1% FA in acetonitrile (solvent B), centrifuged at 800 g for 1 min, soaked in 1-propanol for 1–2 min, and equilibrated with 20 µL of 0.1% FA in water (solvent A). Samples (30 µL) were loaded, taking care to avoid aspirating residual gel debris (when present), centrifuged at 800 g for 1 min, washed with 20 µL of solvent A, and finally overlaid with 150 µL of solvent A before a brief spin to prevent drying.

### Mass Spectrometry

Mass spectrometry was performed on a timsTOF Ultra 2 (Bruker Daltonics, Germany) coupled to an Evosep One system (Evosep Biosystems, Denmark). Peptides were separated using the WhisperZoom 40 SPD gradient on an Aurora Elite 15 × 75 µm, 1.7 µm column (IonOpticks, Australia) with an integrated emitter. The timsTOF Ultra 2 operated in dia-PASEF mode, acquiring full MS scans from m/z 100–1700 and ion mobility 1.45–0.64 1/K₀. The dia-PASEF method covered m/z 400–1000 and 0.64–1.45 1/K₀ with 100 ms accumulation and ramp times. Collision energy was linearly ramped from 20 eV at 0.6 1/K₀ to 59 eV at 1.6 1/K₀. High-sensitivity mode was enabled, and dia-PASEF denoising was disabled.

### Data Processing

DIA data were analyzed using DIA-NN v1.8.1. *In-silico* predicted spectral libraries were generated from a FASTA database comprising the reviewed human proteome (UniProtKB/Swiss-Prot, UP000005640) supplemented with common laboratory contaminants. Search parameters included trypsin digestion with one missed cleavage, variable methionine oxidation and N-terminal acetylation, and N-terminal methionine excision. Precursor charges of 2–4, m/z ranges of 300–1800 (precursor) and 200–1800 (fragment), and mass accuracies of 20 ppm (MS2) and 15 ppm (MS1) were applied. Precursor and protein group FDR thresholds were set to 1%. DIA-NN applied match-between-runs, transferring peptide identification confidence across runs analyzed in the same search to improve quantitative completeness at low input. One-pot samples were processed in a search separate from the baseline and optimal condition samples, precluding identification transfer between one-pot and single-cell data. Baseline and optimal condition samples spanning the tested cell-load range were processed together in the same search. Co-analysis of samples spanning a range of input amounts under match-between-runs maintains low false discovery and false transfer rates even at input differences of two orders of magnitude,^27^ which is beyond the ten-fold cell-load range tested here.

### Statistical Modeling and Functional Enrichment Analysis

Peptide-level quantifications were modeled using the scplainer linear modeling framework implemented in the scp R/Bioconductor package.^28^ A linear model was fit independently for each peptide, with the annotated sample preparation condition and median sample intensity included as covariates to account for differences in total loading amount across runs. Peptides with an insufficient number of observations relative to the number of model parameters were excluded prior to fitting, following scplainer defaults (minimum n/p = 1). The resulting fitted effects and residuals were used to generate batch-corrected and variance-partitioned representations of the data for downstream visualization and comparison across conditions.

To identify functional differences in the proteins recovered under each condition, enrichment analysis was performed using gprofiler2,^29^ querying the Gene Ontology Biological Process, Molecular Function, and Cellular Component terms together with the KEGG and Reactome pathway databases against the *Homo sapiens* genome. Enrichment significance was assessed using the g:SCS multiple-testing correction (threshold = 0.05).^30^ Proteins annotated with transmembrane domains were identified via the UniProt feature field ft_transmem, querying for any annotated transmembrane region. Sequence coverage for transmembrane proteins was calculated as the percentage of residues spanned by uniquely identified peptides relative to the full UniProt protein sequence length. Hydrophobicity of identified proteins was assessed using the Grand Average of Hydropathy (GRAVY) score, calculated as the mean Kyte-Doolittle hydropathy value across all residues in each protein sequence.^31^ Molecular weight distributions were obtained from UniProt-annotated protein mass and compared across conditions in 5 kDa bins.

## RESULTS AND DISCUSSION

### Single-cell images are indexed to proteomic measurements

Same-cell pairing of widefield imaging and discovery-scale LC-MS/MS proteomics uses arrayed thin PAG films, which we term parcels and treat as multi-compartment hydrogel “droplets” that are planar, not spherical (width × length = 2 mm × 5 mm). Each parcel contains an embedded, open top 60-µm diameter microwell for (i) spatially controlled cell isolation and imaging; (ii) simultaneous cell lysis, electrophoresis, and fixation; and (iii) selective removal of hydrogel-encapsulated cell lysates for off-chip trypsin/Lys-C digestion and peptide extraction. Each parcel is spatially addressable to an individual cell or group of cells within the array by standard row-column indexing (Figure 1A-B). A well-characterized human breast epithelial line (MCF-7), provides comparatively large cells (∼19-34 µm)^32,33^ and therefore ample, reproducible protein input for characterizing the methodology. Single and small pools of MCF-7 cells (1-10 cells) are dispensed into the microwells of a dehydrated parcel array by piezoelectric droplet actuation using a single-cell dispenser (cellenONE). The device is then mounted on a widefield microscope for brightfield imaging of each microwell prior to device rehydration and cell lysis. Because each microwell occupies an invariant position in the array, the spatial address of an imaged cell is preserved through all downstream processing, from lysis and electro-injection through parcel excision, in-gel digestion, and LC-MS/MS analysis, allowing the morphological image of each cell to be indexed to the proteomic dataset recovered from that parcel. The 18 × 4 parcel array is designed for parallel processing of up to 72 samples per device. Representative brightfield micrographs of isolated, individual cells paired with total ion chromatograms confirm successful peptide recovery and chromatographic separation from single parcels across independent lanes (Figure 1C), demonstrating end-to-end sample traceability from cell isolation through proteomic measurement.

**Figure 1.**
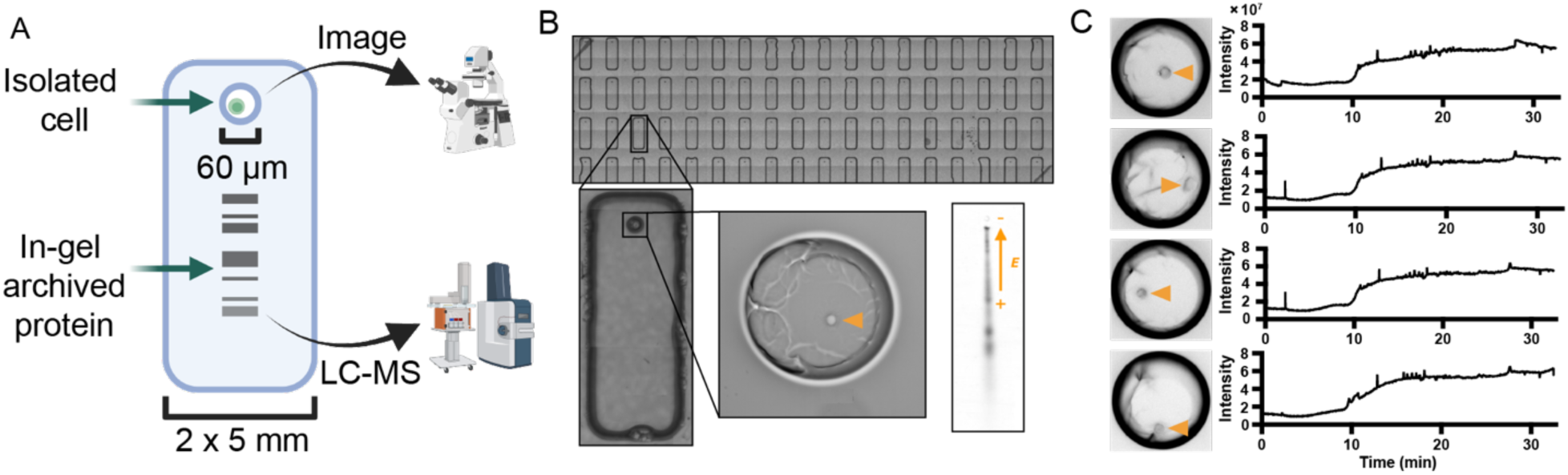
ProteoParcel joint analysis generates imaging and proteomic profiles from the same cell. (A) Schematic of ProteoParcel design and measurement modalities. Single to multiple cells are isolated in 60-µm diameter microwells for imaging prior to lysis and electro-injection into the adjacent PAG. In-gel proteins are digested and extracted for LC-MS/MS. (B) Brightfield micrographs of the parcel array, a single parcel, a single-cell in the patterned microwell, and a representative fluorescence micrograph of stained total protein from a cell lysate. Orange arrow depicts the electric field direction inducing antiparallel electrophoresis. (C) Representative brightfield images of intact cells with their corresponding total ion chromatograms. Orange circles highlight cell position in the microwell.

### Lysis and electro-injection are optimized for proteome solubilization and confinement

We sought to determine the optimal lysis, electrophoresis, and fixation conditions for protein isolation. To do so, we established and characterized three SCP sample preparation configurations. We refer to the conditions as the “baseline condition”, the “optimal condition”, and the “one-pot condition” throughout. Differentiating factors for the baseline and optimal gel conditions are reported in Table 1. Three design rules guided our optimization process; namely, achieving: (i) sufficient solubilization of protein lysate for an effective subsequent peptide digestion, (ii) uniformity of ionic strength across the parcel to reduce electro-injection dispersion, and (iii) capture of each single-cell lysate fully within the bounds of the parcel so that the entire proteome is transferred to LC-MS/MS analysis. Under baseline conditions (0.5% SDS, 15 s electrophoresis, dehydrated gel), SYPRO™ Ruby total-protein fluorescence micrographs revealed diffuse, non-uniform protein bands with a mean axial dispersion, defined as the full-width at half-maximum (FWHM) of horizontal band electropherograms, of 106.2 ± 7.2 µm (n = 4 parcels; Figure 2A-B, Figure S1). We considered two mechanisms for dispersion. First, incomplete solubilization promotes protein aggregation at the microwell interface, reducing the accessible tryptic cleavage sites for trypsin/Lys-C digestion and lowering peptide extraction yield,^34^ which compounds the already limited analyte (∼100-200 pg) available from single mammalian cells.^35^ Second, introducing concentrated lysis buffer into a dehydrated gel (necessary for cell dispensing directly to the bottom of the microwell) for only 30 s generates a transient ionic-strength gradient between the high-conductivity buffer zone in the microwell and the surrounding low-conductivity gel matrix. Because local electric field strength varies inversely with local conductivity, this discontinuity elevates the field in the gel immediately adjacent to the microwell, driving faster, less spatially controlled migration and axial band broadening.^36^ These mechanisms of dispersion were minimized by adjusting the electro-injection parameters.

**Figure 2.**
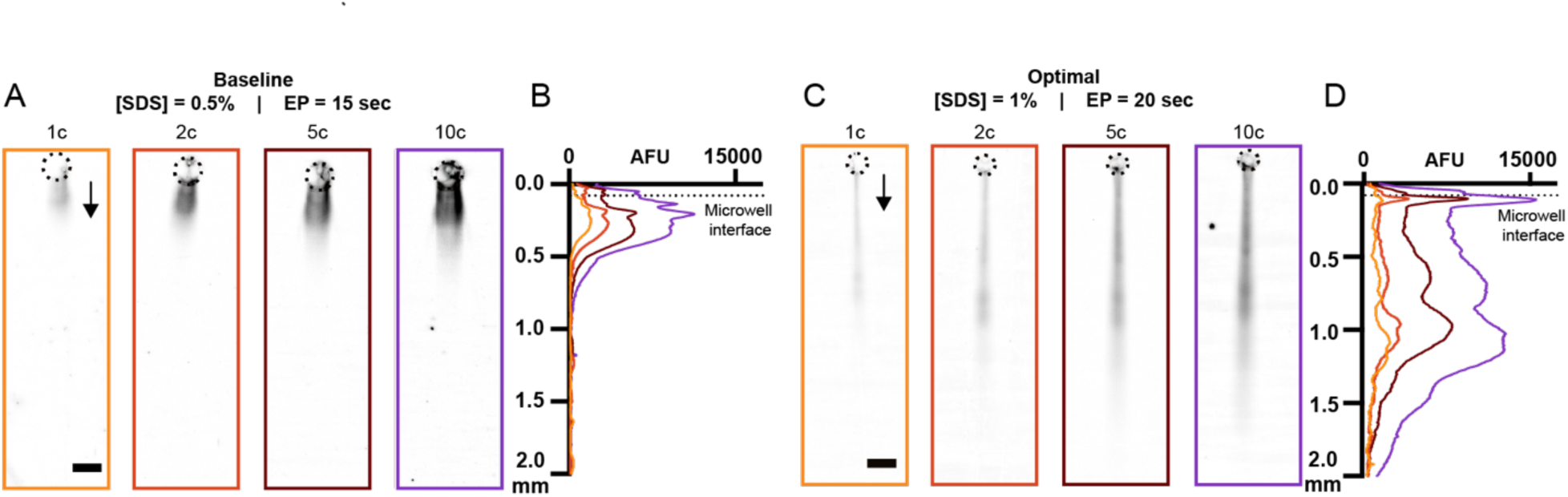
Optimization of lysis and electrophoresis conditions improves electro-injection and reduces band dispersion. (A) Fluorescence micrographs of SYPRO Ruby-stained total protein from 1, 2, 5, and 10 cells after lysis and gel electrophoresis under baseline conditions (0.5% SDS, 15 s electrophoresis). Dashed circles indicate the microwell position. Scale bar = 200 µm. (B) Electropherograms corresponding to the micrographs in (A), showing fluorescence intensity (AFU) as a function of migration distance (mm) for 1 (orange), 2 (dark red), 5 (red), and 10 (purple) cell inputs. The dashed line denotes the microwell– polyacrylamide gel interface; migration proceeds downward. (C) Fluorescence micrographs of SYPRO Ruby-stained total protein from 1, 2, 5, and 10 cells after lysis and gel electrophoresis under optimal conditions (1% SDS, 20 s electrophoresis). Dashed circles indicate the microwell position. (D) Electropherograms corresponding to the micrographs in (C), presented as in (B). Increased SDS concentration and extended electrophoresis duration in optimal result in improved protein injection into the polyacrylamide gel, tighter band migration, and reduced axial dispersion relative to baseline.

**Table 1.** Parcel lysate-encapsulation conditions optimized for LC-MS/MS and proteome depth.

|  | Microwell Diameter | Gel Equilibration | RIPA Buffer Composition | Electrophoresis Time | Digestion Time |
| --- | --- | --- | --- | --- | --- |
| **Baseline Condition** | 100 $\mu\text{m}$ | None (dehydrated) | 0.5% SDS<br>0.25% NaDOC<br>0.5× Tris-glycine<br>0.1% Triton<sup>TM</sup> X-100 | 15 s | 1 h |
| **Optimal Condition** | 60 $\mu\text{m}$ | 5 min, PBS | 1% SDS<br>0.5% NaDOC<br>0.5× Tris-glycine<br>0.2% Triton<sup>TM</sup> X-100 | 20 s | 2 h |

In the optimal condition (1% SDS, 20 s electrophoresis, PBS pre-equilibration), higher SDS improves solubilization, pre-equilibrating the parcel in PBS for 5 min before lysis-buffer addition establishes a more uniform conductivity baseline (a constant diffusion gradient forms from the hydrated gel surface to the underlying glass substrate over the 30 s lysis period), and the longer electrophoresis drives more complete injection. These changes reduced mean band dispersion to 57.0 ± 7.6 µm (n = 4; Figure 2C-D, Figure S2), consistent with more complete proteome solubilization and more uniform electro-injection. Because SDS concentration, electrophoresis duration, and gel hydration state were co-varied between the baseline and optimal conditions, the contribution of each parameter to the improvement in band confinement remains to be independently assessed.

Based on pilot analyses and prior work, the parcel length (5 mm) should accommodate electro-injection and capture of the wide dynamic range of protein molecular masses expected in each cell to ensure that the full proteome is encapsulated in the parcel for downstream digestion and LC-MS/MS.^37^ As such, our efforts to determine the appropriate electrophoresis conditions were further motivated by containing low molecular mass proteins within the lower boundary of the parcel while injecting high molecular mass proteins into the gel contiguous with the microwell interface. The maximum migration distance of the lowest-mass proteins detected by staining is < 2 mm under the optimal condition, well within the 4.22 mm injection lane spanning from the microwell interface to the terminus of the parcel, confirming that each single-cell lysate is encapsulated within the parcel.

### Acid-alcohol fixation immobilizes proteins in the parcel

In-gel digestion requires that injected proteins remain encapsulated in the parcel through electro-injection and washing while remaining reactive to trypsin/Lys-C and extractable as peptides after in-gel digestion. The scWB assay that inspired the form factor of the parcels introduced in this work immobilizes proteins covalently to a PAG layer by co-polymerizing N-[3-[4-(benzoylphenyl)formamido]propyl]methacrylamide (BPMAC) into the PAG network during gel polymerization for subsequent UV-initiated protein-target capture to said light-reactive PAG polymer network.^21^ We hypothesized that this covalent linkage to PAG via BPMAC would sterically occlude trypsin/Lys-C cleavage sites on immobilized proteins and physically retain tryptic peptides within the matrix after digestion, causing peptide loss during osmotic extraction and, thus, reduced LC-MS/MS yield. Supporting this, *in situ* measurements of thermodynamic partitioning show that UV photoactivation of BPMAC increases the equilibrium partition coefficient of model proteins toward the gel phase relative to unmodified PAG (p = 0.0286, Mann-Whitney U-test)^38^, a property well-suited to immunoassays but counterproductive for workflows requiring downstream peptide extraction from a PAG scaffold.

We therefore tested acid-alcohol precipitation as a fixation approach. Acid-alcohol precipitation would encapsulate proteins in the hydrogel parcel without forming covalent protein-polymer bonds, while preserving cleavage-site accessibility, peptide release, and residual SDS (a downstream MS interferent) washout.^39,40^ To test whether covalent protein-polymer crosslinking restricts protein accessibility relative to non-covalent acid-alcohol precipitation, we performed in-gel native immunoprobing of GFP-expressing MCF-7 cell lysates to compare antibody-based detection of GFP and beta-tubulin for lysate fixed under each chemistry. Acid-alcohol fixation resolved three distinct immunoreactive species (beta-tubulin, GFP, and a GFP isoform), whereas BPMAC fixation recovered primarily one GFP species under equivalent probing parameters (Figure S3). The reduced species detection under BPMAC suggests either covalent modification thus reducing protein accessibility within the gel matrix or incomplete photocapture thus allowing diffusive protein loss during processing. Based on these results, we adopted acid-alcohol as the fixation chemistry for downstream in-gel digestion and LC-MS/MS.

### Proteome quantification and sensitivity

Having selected lysis and electrophoresis parameters that improve solubilization and proteome encapsulation, we next evaluated the downstream effect of each preparation method on single-cell proteome depth (Figure 3). We set digestion conditions by measuring residual in-gel protein signal after digestion and osmotic extraction across enzyme amount (2 or 5 ng trypsin/Lys-C) and time (1-2 h at 70 °C) (Figure S4). Peptide extraction efficiency between 2 and 5 ng of enzyme was quantified as the percentage loss of total protein staining signal after digestion. Extraction efficiency improved most at higher cell loads (10 cell samples: +31% from 2 to 5 ng of trypsin/Lys-C) and, interestingly, decreased at the single-cell level (-15.5% from 2 to 5 ng of trypsin/Lys-C). A recent study evaluating key parameters in SCP sample preparation indicates minimal improvement in protein coverage completeness and peptide detection between 1-10 ng of trypsin and a negative correlation between trypsin amount and proteins identified.^11^ As such, we applied a 2-h digestion with 2 ng of trypsin/Lys-C for the optimal condition compared to the 1-h digestion with 2 ng of enzyme for the baseline condition.

**Figure 3.**
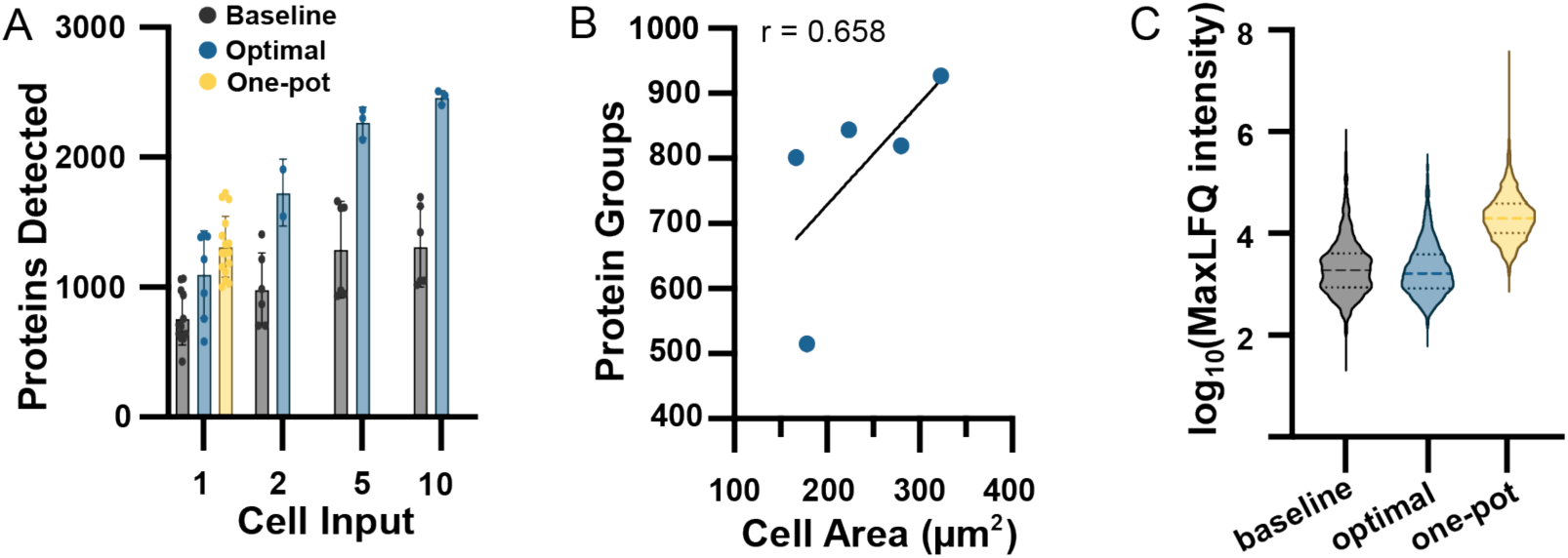
ProteoParcel pairs morphology with proteome measurement. (A) Number of detected protein forms across varying cell loads for baseline (gray), optimal (blue) and one-pot (yellow) conditions. (B) Brightfield cell cross-sectional area versus protein-group identifications for n = 5 imaged single cells (optimal); each point is one cell, line is the linear regression (r = 0.658, p = 0.228). (C) Dynamic range of protein intensities detected in single-cell samples.

We next sought to evaluate the quantitative depth of each sample preparation method. Protein identifications were compared across baseline, optimal, and an in-solution one-pot control. Under this latter one-pot condition cells are dispensed, lysed, and digested directly in 1 µL of rapid digestion mix (2 ng of enzyme total). Baseline and optimal conditions were also assessed across small cell pools of 2, 5, and 10 cells per microwell. After batch correction and cell filtering using the scplainer pipeline, the optimal condition returned an average of 1098 ± 337 protein IDs per single cell (n = 7), and the baseline condition returned an average of 760 ± 203 (n = 12) (Figure 3A). One-pot digestion samples returned 1312 ± 233 protein IDs per single cell (n = 16), establishing an upper benchmark for gel-free preparation under equivalent LC-

MS/MS conditions. Median peptides identified per cell followed the same trend (baseline condition: 1949 peptide IDs, optimal condition: 4027 peptide IDs for optimal, and one-pot condition: 4383 peptide IDs; Figure S5). Protein identifications scaled with the number of cells pooled for both gel conditions, confirming that increased analyte load is reflected in deeper proteome coverage (Figure 3A).

We sought to index single-cell morphology to proteome depth for selected cells. Across the imaged single cells, larger cells trended toward more protein-group identifications (r = 0.658, p = 0.228, n = 5; Figure 3B). Unnormalized protein intensity values trend positively with cell size (r = 0.71, p = 0.18) as well as unnormalized precursor intensity values (r = 0.68, p = 0.21)(Figure S6). While the trends did not reach statistical significance (likely due to low sample size), single-cell proteomic depth is known to correlate with cell size.^41^ Observation of cell morphology and identified proteins from the same cell demonstrate image-to-proteome integration.

We next wanted to determine the dynamic range of protein quantification. Single-cell samples spanned 4.3, 3.3, and 4.4 orders of magnitude in protein intensity for the baseline, optimal, and one-pot conditions, respectively (log_10_-transformed MaxLFQ intensities; Figure 3C).The unexpectedly wide range in the baseline gel condition reflects identification of proteins supported by a single cell, where the reported intensity is a single, unreplicated measurement rather than a reproducible signal: 182 of 1027 proteins (17.7%) in the baseline condition were detected in only one cell. These singly-detected proteins disproportionately populate the low end of the intensity distribution. Restricting to proteins detected in at least two cells decreases the dynamic ranges to 3.4, 2.8, and 4.2 orders of magnitude (-0.9, -0.5, and -0.2) for the baseline, optimal, and one-pot conditions, respectively. These more stringently determined values suggest the true dynamic range for parcels is ∼3 orders of magnitude.

### Single-cell reproducibility and variance analysis

Quantitative reproducibility, assessed as the coefficient of variation (CV) of log_2_-transformed protein intensities across single-cell replicates, was low in all conditions, with median CVs of 0.084 (baseline condition, n = 1094 protein IDs), 0.073 (optimal, n = 1224 protein IDs), and 0.062 (one-pot, n = 1901 protein IDs) (Figure 4A). The comparable CV values across gel-based and in-solution conditions indicate that parcel preparation does not substantially increase quantitative variability, and the modest monotonic decrease observed from baseline to optimal to one-pot conditions is consistent with incremental gains in protein solubilization and extraction completeness.

**Figure 4.**
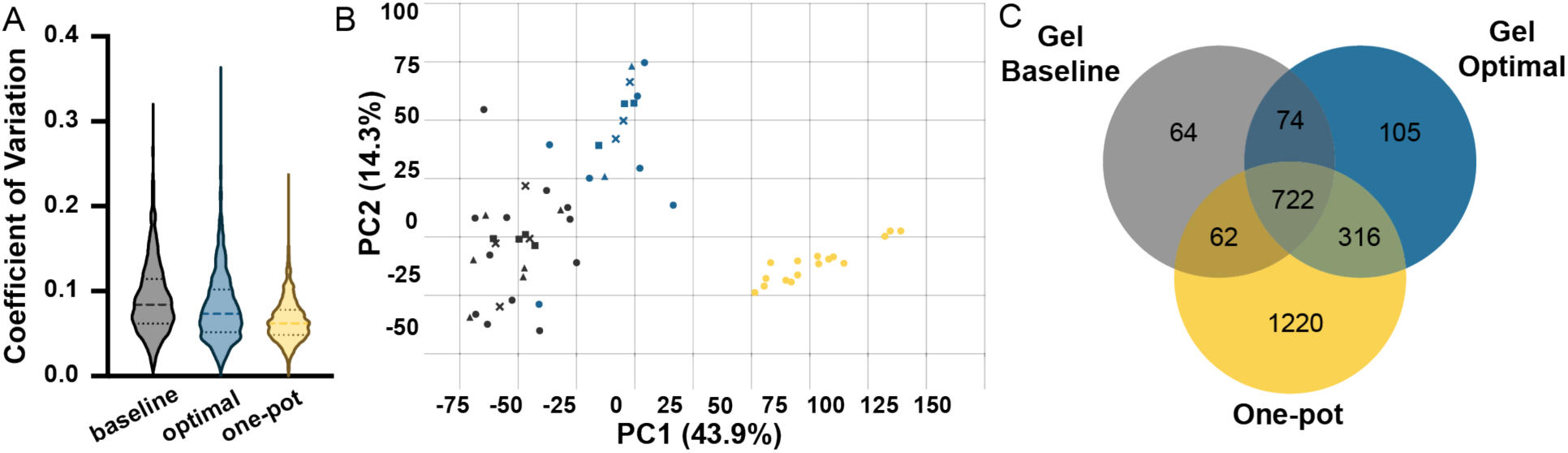
Quantitative reproducibility and proteome composition across ProteoParcel conditions (A) Coefficients of variation of log₂-transformed protein intensities; the dashed line is the median. (B) APCA+ dimensionality reduction on the sample preparation condition effect. 1-cell: circles; 2-cell: triangles; 5-cell: squares; 10-cell: crosses. (C) Venn diagram of uniquely identified proteins in each sample preparation method. Unique proteins are subsampled to 7 runs per condition for 100 iterations.

To quantify the sources of proteome-level variance across conditions, we applied the scplainer linear modeling framework (see Experimental Section). Variance partitioning attributed ∼31% of peptide-level and ∼33% of protein-level variance to per-cell loading amount (Median Intensity), and ∼22% of peptide-level and ∼19% of protein-level variance to sample-preparation condition (Figure S7). The remainder was attributable to unexplained residual variation, consistent with the high cell-to-cell heterogeneity characteristic of single-cell proteomics datasets.^28^ To isolate the condition-specific effect from loading variation, we applied augmented principal component analysis (APCA+) with sample-preparation condition as the effect of interest (Figure 4B).^42^ PC1 (43.9% of variance in the condition subspace) separated one-pot from both gel conditions. PC2 (14.3%) resolved the baseline from optimal conditions, with no sub-clustering by cell input within any condition. Variance partitioning quantified a distinct share of variance attributable to sample preparation condition, and APCA+ analysis revealed distinct clustering by sample preparation method. Together, these results confirm that the preparation method exerts a systematic, quantifiable effect on the recovered proteome, separate from total protein input, across the 1–10 cell range tested.

To compare the proteins identified across conditions, unique identifications were subsampled to seven runs per condition over 100 iterations. All three conditions identified 722 proteins in common, representing the core detectable MCF-7 proteome across sample preparation methods (Figure 4C). The optimal condition identified 316 protein forms in common with one-pot, increased from the 62 shared between the baseline and one-pot conditions, which is consistent with improved solubilization, injection, and peptide extraction of the optimal condition over the baseline condition. One-pot uniquely identified 1,220 protein forms detected in neither parcel condition, motivating continued optimization of in-parcel digestion and peptide extraction for future ProteoParcel workflows.

### Parcel preparation broadly preserves functional and physiochemical diversity

We sought to understand whether the ProteoParcel workflow introduces specific protein-class loss. To do this, we scrutinized eight different categories of typical mammalian-cell housekeeping proteins. The categories were histones, proteasome subunits, mitochondrial electron transport chain components, metabolic enzymes, translation factors, chaperones, cytoskeletal proteins, and ribosomal proteins. The baseline condition identified 353 proteins, the optimal condition identified 384 proteins, and the one-pot condition as a benchmark identified 534 proteins in these eight categories (Figure 5A). All three conditions show similar relative proportions between the annotated categories despite different totals indicating that the proteins uniquely identified in each condition are likely not drawn from a narrow functional niche.

**Figure 5.**
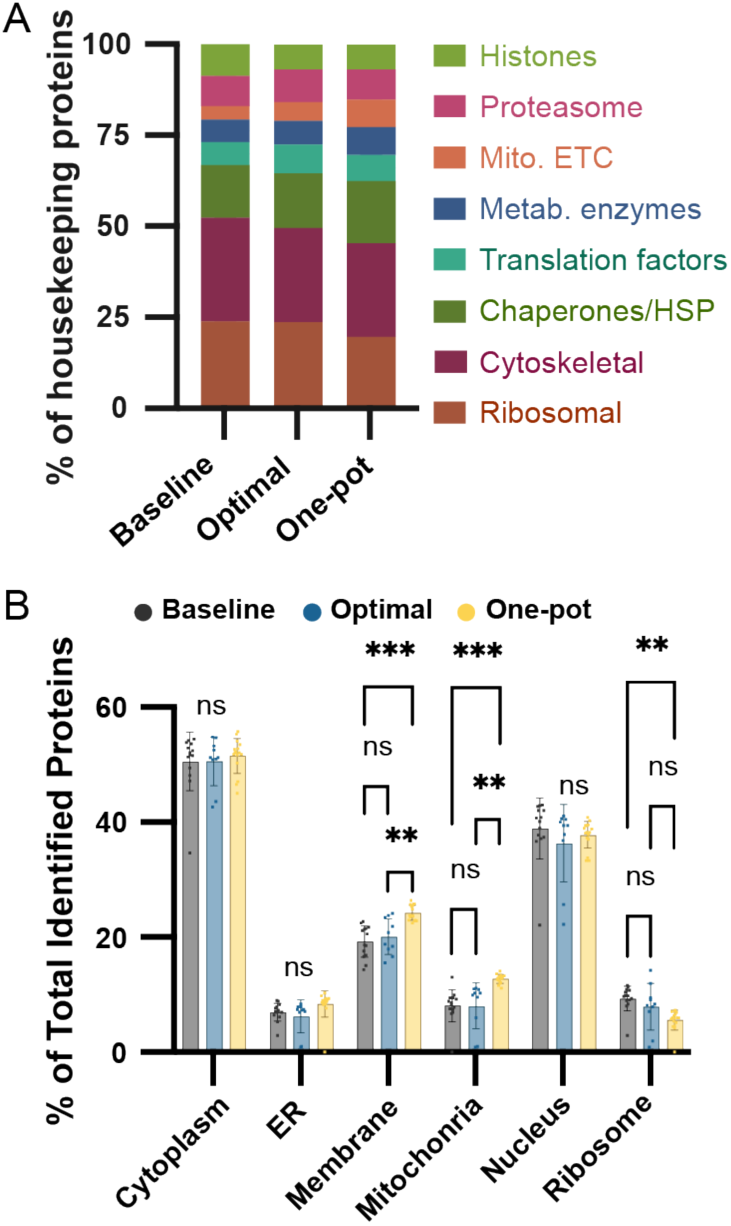
Single-cell proteomes analyzed by LC-MS/MS show minimal bias toward functional groups and subcellular compartments. (A) Stacked bar plot showing functional protein class composition for baseline (n = 353), optimal (n = 384), and one-pot (n = 534) conditions across eight housekeeping categories. Proteins unannotated to these categories (∼70% per condition) are excluded. (B) Grouped bar plot showing the percentage of identified proteins localizing to six subcellular compartments for baseline (gray), optimal (blue), and one-pot (yellow). Two-way ANOVA with Tukey’s post-hoc test (ns, not significant; **p < 0.01; ***p < 0.001).

Cytoplasm and nucleus were the most represented subcellular compartments across all three conditions, each accounting for roughly 50% and 37–39% of identified proteins, respectively, consistent with the expected abundance distribution of a mammalian cell line proteome (Figure 5B).^43^ In contrast, membrane and mitochondrial compartments were significantly depleted in both parcel conditions relative to the one-pot condition (two-way ANOVA with Tukey post-hoc; baseline vs one-pot p < 0.001; optimal vs one-pot p < 0.01; Figure 5B). This pattern is consistent with the known difficulty of extracting long hydrophobic peptides from acrylamide gels.^44,45^ Specifically, integral membrane proteins stabilized by an SDS-micelle coating can aggregate within the gel pores decreasing solubilization during peptide extraction.^46^ Depletion of membrane proteins was slightly attenuated under the optimal condition relative to the baseline (mean detection: baseline condition = 19.3%, optimal condition = 20.1%), indicating that more complete lysis and longer digestion improve hydrophobic protein recovery. However, the reduced number of identified membrane proteins persisted under the optimal condition relative to the one-pot condition. This loss points to peptide extraction from the gel matrix, rather than incomplete lysis, as the dominant peptide-loss mechanism. Beyond peptide loss characterization, consistent coverage across cellular compartments is a prerequisite for relating paired subcellular imaging to proteome depth.

Transmembrane proteins were detected less by both identifications and intensity in the baseline and optimal conditions, and the transmembrane proteins that were detected were covered by fewer peptides per protein as compared to the one-pot condition (Figure 6A). For the 10 transmembrane proteins detected across all conditions with the highest unique peptide counts, the one-pot condition achieved the highest per-protein sequence coverage, with a mean overall coverage of 30.3% compared to 20.0% (optimal) and 17.7% (baseline). This observation is consistent with solution-phase digestion providing trypsin unobstructed access to the hydrophilic extracellular and cytoplasmic loops flanking transmembrane helices. The optimal condition exceeded the baseline condition in eight of the top ten proteins, a modest gain attributable to better solubilization of transmembrane domains and increased enzyme diffusion during digestion in the optimal versus baseline condition. In-gel digestion proceeds by diffusion of trypsin and Lys-C through the PAG matrix rather than bulk transport, so peptide yield increases with extended incubation time.^47,48^

**Figure 6.**
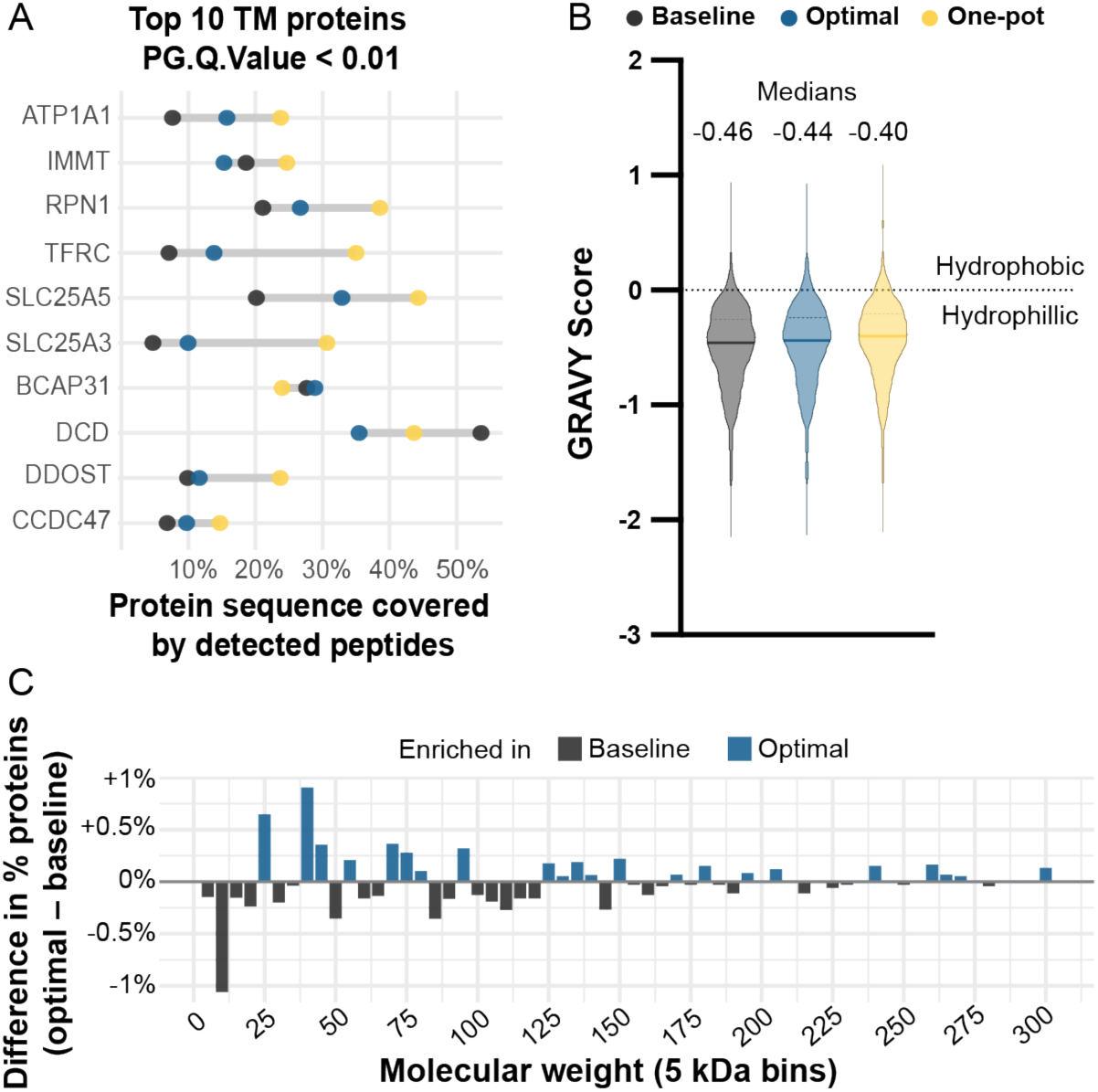
Physicochemical characterization of detected proteomes across sample preparation conditions. (A) Dot plot showing whole-sequence coverage by detected peptides for the top 10 transmembrane proteins per condition. Each dot represents mean sequence coverage per protein per condition. (B) Violin plots showing the distribution of Grand Average of Hydropathy (GRAVY) scores per condition. Medians indicated above. The dashed line at y = 0 denotes the hydrophobic/hydrophilic boundary. (C) Difference in protein detection gel conditions across molecular weight bins (5 kDa), expressed as the difference in percentage of total identified proteins (optimal – baseline conditions).

When considering whole-proteome readout of hydrophobicity bias, the distribution of Grand Average of Hydropathy (GRAVY) scores across all detected proteins provides an independent way to assess that bias (Figure 6B).^31^ Protein GRAVY score distributions were centered below zero in all three conditions (medians: baseline = -0.46; optimal = -0.44; one-pot = -0.40), confirming that hydrophilic proteins dominate the detected proteome regardless of preparation method as shown in previous studies.^49^ The medians shift monotonically toward less-negative values, from -0.46 to -0.44 to -0.40. This shift traces slightly higher hydrophobic protein coverage as extraction constraints relax. The signal is small but resolved, and the shift corroborates the compartment and transmembrane protein coverage findings through an independent physicochemical axis. The direction of this shift follows from the mechanics of in-gel peptide recovery. Peptide extraction from the gel is itself diffusion-limited and peptide-dependent, with recovery decreasing for larger, more hydrophobic peptides that exhibit non-specific adsorption to the PAG backbone.^48^ Peptide-level hydrophobicity analysis showed a similar trend of more hydrophobic peptides detected in one-pot samples (median GRAVY = - 0.18) with significant improvement in detection of hydrophobic peptides in optimal condition (median GRAVY = -0.41) compared to baseline conditions (median GRAVY = -0.49) (Figure S8). Sequential extraction with ACN drives osmotic shrinkage of the gel matrix, improving peptide release and collection prior to LC-MS/MS.

We next analyzed the molecular mass distribution of detected proteins to identify any underlying mass-dependent shifts in detection between gel conditions. Figure 6C plots the percent difference in protein identifications (optimal – baseline) across 5-kDa molecular mass bins. Relative to baseline, the optimal condition was enriched for proteins in the 20-45 kDa range. The largest single-bin gains fell at 25-30 kDa (+0.65%) and 40-45 kDa (+0.91%), with cumulative enrichment of +1.60% across all bins ≥25 kDa. Conversely, the baseline condition showed a corresponding excess of low-mass proteins, most notably below 15 kDa (-1.06% at 10-15 kDa; -0.94% overall for proteins <30 kDa). Differences above 50 kDa were small (e.g., +1.36% cumulative ≥120 kDa but -0.21% cumulative ≥100 kDa) and did not follow a consistent direction, indicating that the gain in the optimal condition is concentrated in the mid-mass range rather than reflecting a uniform shift toward higher-mass species. The near-identical median masses (baseline 47.1 kDa; optimal 46.9 kDa) confirm that the overall distribution is not grossly shifted. The bin-level pattern instead reflects a targeted gain in mid-mass protein recovery and a corresponding reduction in low-mass protein representation under the optimal condition.

During electro-injection, the 7%T PAG matrix acts as a molecular sieve, slowing proteins with hydrodynamic radii approaching the gel pore size. Proteins of higher mass or poor solubility may aggregate at the microwell interface, reducing the efficacy of enzymatic digestion compared to lower mass species. During digestion, diffusion-limited enzyme access and peptide recovery further disfavor larger species.^48^ Higher SDS concentration and extended digestion time in the optimal condition partially compensate, reducing aggregation of hydrophobic species before fixation and allowing more complete proteolysis: median unique peptides per protein were 2, 3, and 2 for baseline, optimal, and one-pot conditions, respectively (Figure S9). Additionally, missed cleavage rate (≥1 missed cleavage), determined by counting internal lysine or arginine residues not followed by proline in each unique detected peptide sequence, was 42.1%, 32.0% and 16.4% for baseline, optimal, and one-pot conditions, respectively, further supporting optimization of solubility and proteolysis.

## CONCLUSION

To pair brightfield microscopy of live, intact cells with bottom-up single-cell mass spectrometry proteomics of that same cell we introduce ProteoParcel. To index whole-cell images with that same cell’s proteome, we electrophoretically inject whole-cell lysates into arrayed parcels, drawing each lysate from a microwell holding a single cell. Each parcel occupies a fixed position within the array. That fixed position preserves the spatial index linking each imaged cell to its downstream in-gel digest. Optimization of lysis buffer composition, electrophoresis duration, and gel pre-equilibration conditions reduced mean axial band dispersion by 1.9-fold relative to non-optimal conditions while maintaining proportional protein recovery across 1-10 cell inputs. This is a significant improvement for reducing protein aggregation at the microwell interface which we have hypothesized leads to reduced tryptic digestion efficiency and osmotic peptide extraction. Acid-alcohol fixation was validated as a viable non-covalent alternative to BPMAC photocapture for encapsulation of limited analytes in PAG matrices. Non-covalent fixation preserves downstream trypsin cleavage site accessibility and peptide extractability while intrinsically removing SDS prior to LC-MS/MS analysis.

Through optimization of protein solubilization from cell lysates, electro-injection dispersion, and in-gel digestion, ProteoParcel reports a maximum of >1,400 protein forms from a single-cell in-gel digest. Systematic characterization of single-cell proteome composition across parcel and in-solution methods reveal that PAG matrix interactions introduce predictable physicochemical shifts rather than stochastic or functionally arbitrary losses. Mechanism-consistent phenomena driving proteomic shifts along subcellular localization, hydropathic, and molecular mass axes are revealed. Functional class proportionality represented by key protein classes (e.g., ribosomal proteins, histones, and metabolic enzymes) is broadly preserved and consistent across preparation methods. Membrane and mitochondrial compartment representation, transmembrane protein sequence coverage, GRAVY score distributions, and mass-dependent detection patterns collectively point to SDS solubilization, digestion time, and peptide extraction as the primary determinants of residual bias relative to in-solution digestion of single cells.

ProteoParcel establishes a technical foundation for hydrogel-based sample preparation in single-cell multiomics, leveraging the PAG matrix as a multifunctional planar “droplet” analog for cell isolation, proteome archiving, and MS preparation. Multimodal single-cell measurements deepen our understanding of how biomolecular expression underlies phenotypic heterogeneity in complex samples. By pairing brightfield morphological imaging with discovery-scale proteomics from the same cell, ProteoParcel makes this combination accessible through microscale manipulations and plate-based sample preparation without specialized liquid handling infrastructure. By physically confining intact single-cell proteomes within mesoscale PAG parcels, spatial indexing of each cell’s identity is preserved from the microscope to the mass spectrometer. In characterizing the ProteoParcel workflow, we have also identified and described physicochemical phenomena governing tryptic digestion efficiency and proteome recovery from microfluidic gel substrates. Looking ahead, we anticipate that the ProteoParcel LC-MS/MS workflow can be extended to BPMAC-fixed gels archived from scWB experiments enabling retrospective pairing of targeted, proteoform-resolved immunodetection with unbiased, discovery-based bottom-up proteomics from the same single-cell. This would open a new axis of targeted single-cell multiomics inaccessible to existing platforms. Further, we infer from our demonstrated imaging integration that additional inverted microscopy-based modalities may be applied, such as fluorescent reporter bioassays, cell migration assays, cell interaction measurements, and subcellular resolution microscopy.

