## Supplementary Information for "Arrayed hydrogels pair whole-cell imaging with single-cell mass spectrometry proteomics"

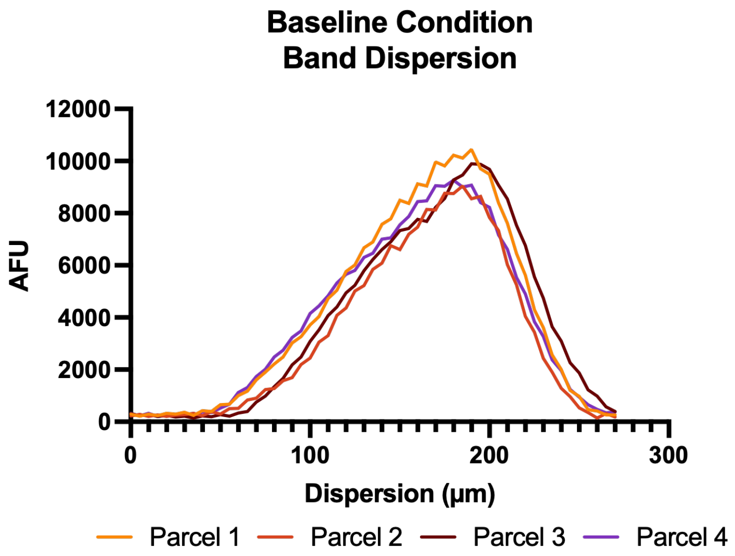


**Supplemental Figure S1**. **Band dispersion profiles for baseline condition.** Horizontal band profiles of stained protein from four single-cell baseline condition parcels depicted in main text Figure 2A (representative micrograph, "1c"). Mean axial dispersion was calculated from these data as the full-width at half-maximum and is reported in the main text.

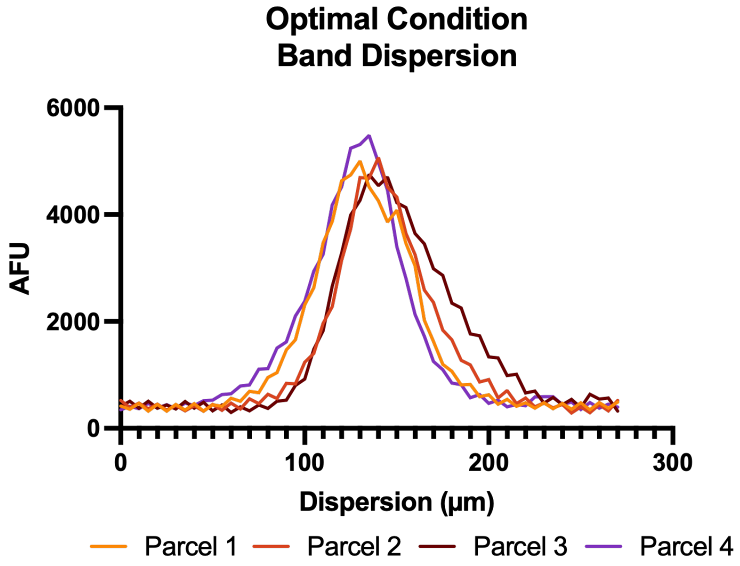


**Supplemental Figure S2**. **Band dispersion profiles for optimal condition.** Horizontal band profiles of stained protein from four single-cell optimal condition parcels depicted in main text Figure 2C (representative micrograph, "1c"). Mean axial dispersion was calculated from these data as the full-width at half-maximum and is reported in the main text.

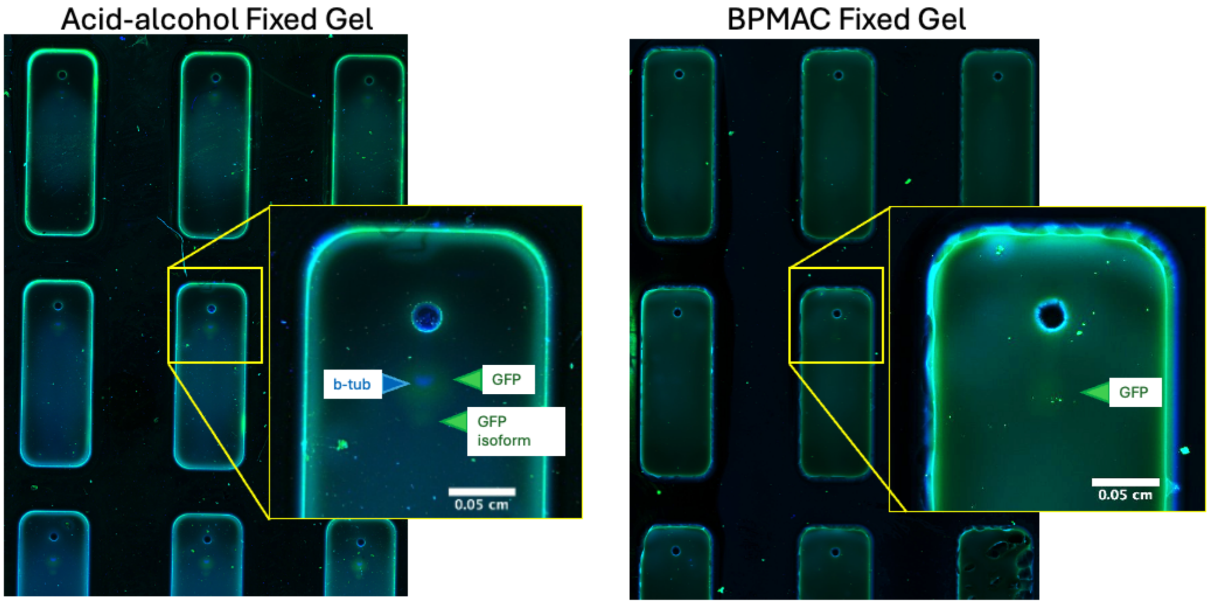


**Supplementary Figure S3. In-gel native immunoprobing of GFP-expressing MCF-7 cell lysates in acid-alcohol-fixed and BPMA-fixed conditions.** Representative micrographs of parcels immunostained for GFP and beta-tubulin under the optimal RIPA buffer and electrophoresis conditions specified in main text Table 1. Under identical immunoprobing conditions, beta-tubulin and two isoforms of GFP were detected in acid-alcohol fixed gels while only one isoform of GFP was resolved in BPMA-fixed gels.

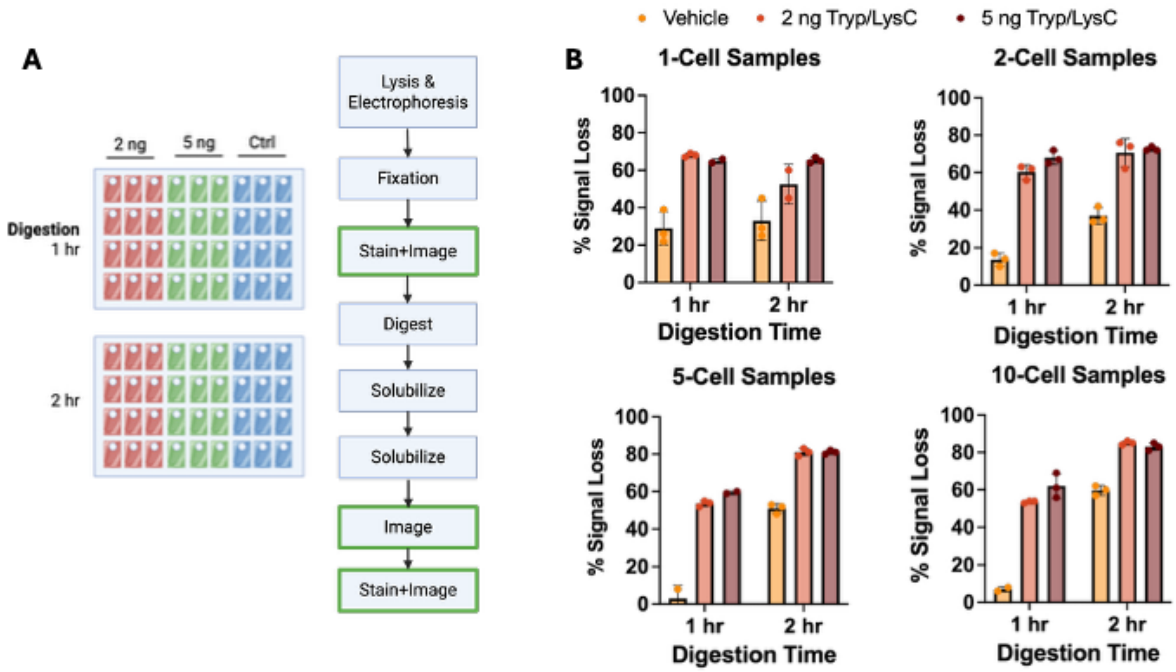


**Supplementary Figure S4. In-gel digestion optimization varying cell load, digestion time, and mass of enzyme.** (A) Experimental configuration for digestion optimization. Two ProteoParcel devices were prepared with subsets of parcels subjected to 2 ng of trypsin/Lys-C, 5 ng of trypsin/Lys-C, or a vehicle control containing no enzyme. (B) Percentage of signal loss measured as the difference between total SYPRO Ruby fluorescence signal before digestion and after digestion and re-staining as a percentage of total signal before digestion.

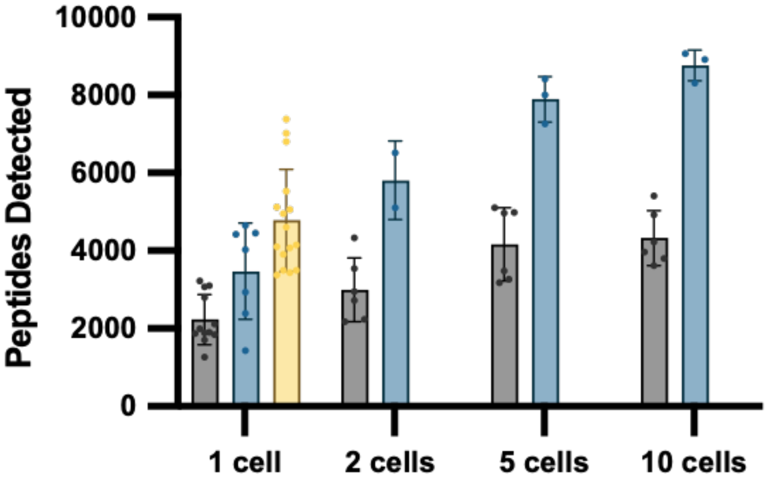


**Supplementary Figure S5. Number of detected peptides across varying cell loads for all sample prep conditions.** Bar plots of unique peptides identified in the baseline condition (grey), optimal condition (blue), and one-pot condition (yellow). As reported in the main text, median peptides identified for single-cell samples are 1949 peptide IDs in the baseline condition, 4027 peptide IDs in the optimal condition, and 4383 peptide IDs in the one-pot condition.

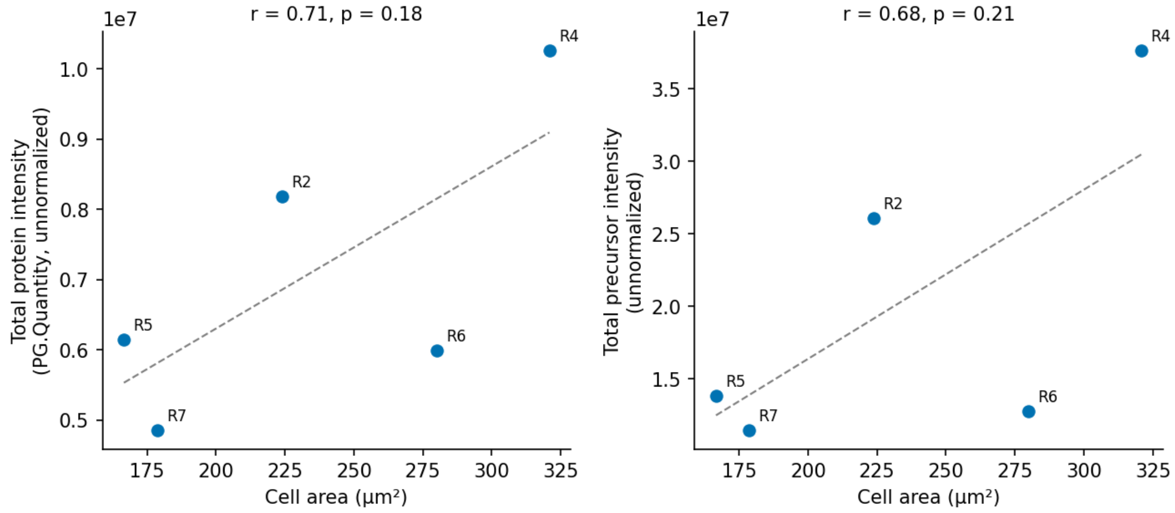


**Supplementary Figure S6. Protein and precursor intensities by cell area for paired MCF7 single-cell replicates.** Scatter plots of imaged cell area ($\mu m^2$) versus (left) total protein intensity (sum of unnormalized PG.Quantity across all identified protein groups) and (right) total precursor intensity (sum of unnormalized Precursor.Quantity) for five single-cell DIA runs (R2, R4, R5, R6, R7; labeled). Dashed lines show linear regression fits.

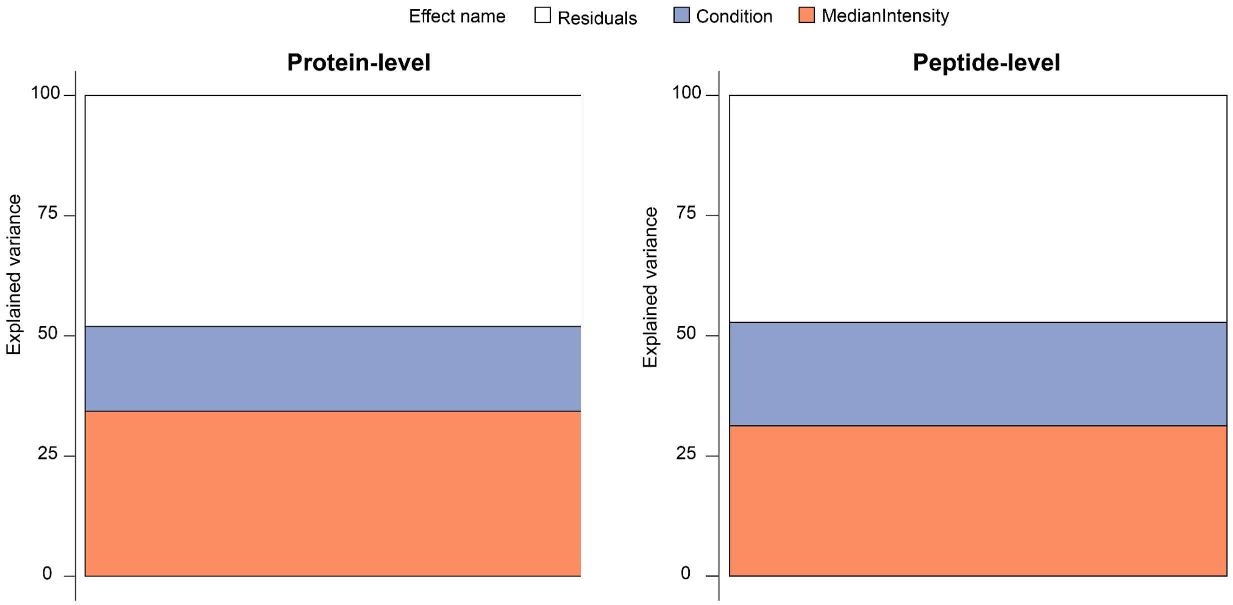


**Supplementary Figure S7. Variance partitioning of protein- and peptide-level abundance across sample-preparation conditions.** Stacked bar plots of scplainer linear model (~ 1 + Condition + MedianIntensity) variance decomposition for all datasets (baseline condition, optimal condition, and one-pot), averaged across all quantified features. (Left) Protein-level variance: per-cell loading amount (MedianIntensity) accounts for ~33% and sample-preparation condition for ~19% of total variance, with the remaining ~48% attributed to residual (unexplained) variation. (Right) Peptide-level variance: MedianIntensity accounts for ~31% and condition for ~22%, with ~47% residual.

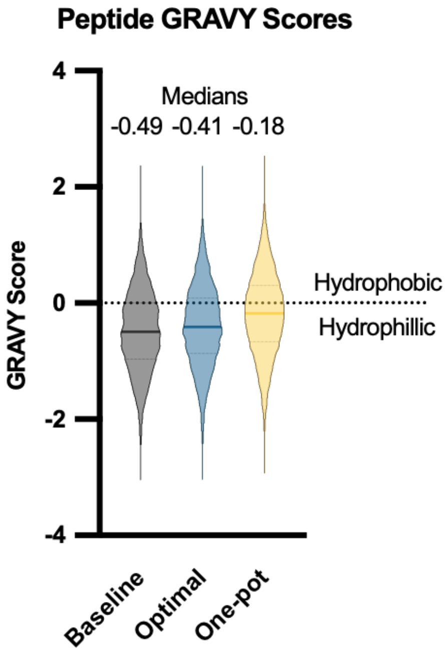


**Supplementary Figure S8. Peptide-level hydrophobicity distribution.** Violin plots showing the distribution of Grand Average of Hydropathy (GRAVY) scores at the peptide-level per condition. Medians indicated above the plots. The dashed line at y = 0 denotes the hydrophobic/hydrophilic boundary.

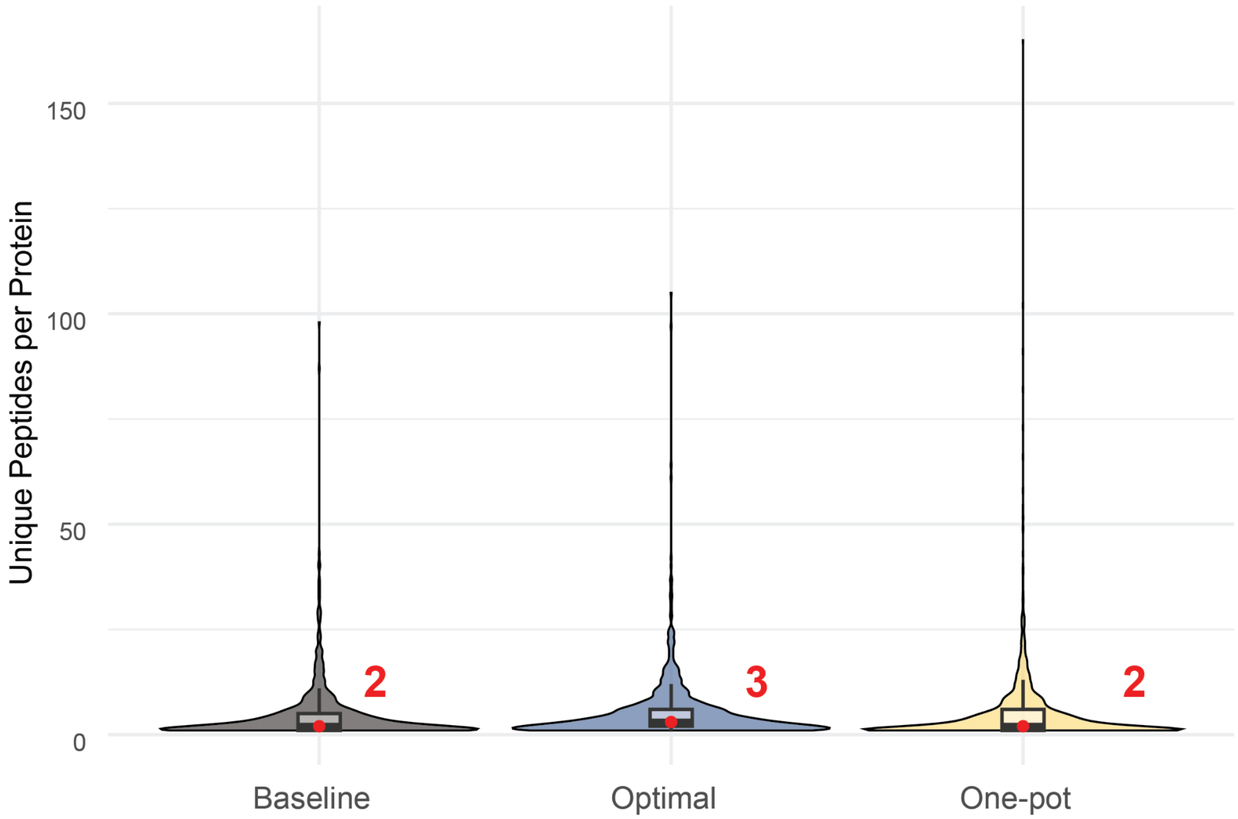


**Supplementary Figure S9. Unique peptides per identified protein across sample-preparation conditions.** Violin and box plots of the number of unique peptide sequences identified per protein for the baseline (grey), optimal (blue), and one-pot (yellow) conditions. Red points and labels indicate condition medians (2, 3, and 2 unique peptides per protein, respectively).
